# Dose-response surface fits to drought and nitrogen limitation applied together allow mapping of loci that exhibit nonlinear responses

**DOI:** 10.1101/186791

**Authors:** Megan M. Chang, Danielle Allery Nail, Toni Kazic, Susan J. Simmons, Ann E. Stapleton

## Abstract

Crop improvement must accelerate to feed an increasing human population in the face of environmental changes. Breeding programs can include anticipated climatic changes and genetic architecture to optimize improvement strategies. We analyzed the genetic architecture underlying the response of Zea mays to combinations of water and nitrogen stresses. Recombinant inbreds were subjected to nine combinations of the two stresses using an optimized response surface design, and their growth was measured. Three-dimensional dose response surfaces were fit globally and to each polymorphic allele to determine which genetic markers were associated with different response surfaces. Three quantitative trait loci that produced nonlinear surfaces were mapped. Alleles that performed better in combinations of mid-range stresses were typically not the alleles that performed best under combinations of extreme stresses. To develop physiologically relevant models for future genetic analyses, we modeled the network that explains the response surfaces. The network contains two components, an elliptical paraboloid and a plane, that each combine the nitrogen and water inputs. The relative weighting of the two components and the inputs is governed by five parameters. We estimated parameter values for the smoothed surfaces from the experimental lines using a set of points that covered the most distinctive regions of the three-dimensional surfaces. Surfaces computed using these values reproduced the smoothed experimental surfaces well, especially in the neighborhood of the peaks, as judged by three different criteria. The parameters exaggerated the amplitudes of the simulated surfaces. Experiments using single stresses could misestimate responses to their combinations and disguise loci that respond nonlinearly. The three-dimensional shape evaluation strategy used here more thoroughly compares nonlinear, nonplanar responses. We encourage the application of our findings and methods to experiments that mix crop protection measures, stresses, or both, on elite and landrace germplasm.

## Introduction

Crop improvement will need to accelerate in the coming decade, as the human population increases and the abiotic environment changes (Wheeler and von Braun 2013). Cultivar improvement in stress resistance is key. While breeding programs can increase the rate of genetic gain by using genotype prediction to shorten the breeding cycle time, (Jonas and de Koning 2013), crop improvement depends on the immediate agricultural context, complicating selection schemes (*Cooper et al.* 2014). So far, cultivar improvement in maize has not significantly increased stress tolerance on a large scale (Lobell *et al.* 2014). But there is ample potential for improvement: the maximum yield in yield competitions is typically one-third higher than the average yield in comparable fields (Tollenaar and Lee 2002).

Crop performance in a variety of agricultural contexts is an example of a complex phenotype. Each context differs in the combinations of extant stresses, and the magnitudes of each individual stress, that the plants experience: and the responses of different genotypes to those contexts can differ considerably. Ensuring the strong phenotypic signal needed for selection requires knowing where the genetic architecture of the phenotype lies on the continua of a few polymorphisms exerting large effects, *vs.* many polymorphisms with small effects, because breeding strategies necessarily differ. Thus, the costs and difficulties of breeding escalate as the numbers of contexts and genotypes, and phenotypic complexity, increase.

Breeding programs can optimize the use of their resources by selecting germplasm in the most predictive environments (Hallauer *et al.* 2010). However, this current-environment approach does not necessarily include the most predictive environments under climate change (Cairns *et al.* 2013; Makumburage *et al.* 2013). To predict performance in the out-of-range field environments of the future, we will need a mechanistic view that incorporates climate prediction, knowledge of the phenotype’s genetic architecture, and understanding of the physiological systems controlling the complex phenotype of response to combined stresses (Cooper *et al.* 2014; Heslot *et al.* 2014).

There are examples of genetic dissection of abiotic stress combinations for the combination of heat and drought, and ultraviolet radiation (UV) plus drought (Cairns *et al.* 2013; Makumburage *et al.* 2013). In the (Cairns *et al.* 2013) study, genotypes with good performance in drought or heat did not perform well in the combined heat and drought environments. This is an example of the breeder’s rule that selection for improvement should be carried out in the target environment (Hallauer *et al.* 2010). A controlled greenhouse-scale analysis of high UV stress combined with moderate drought also indicated that loci important for one stress did not have an important effect in the combined stress treatment; and that the combined stress effect level was less than additive, indicating a nonlinear protective interaction between the two stresses (Makumburage *et al.* 2013).

Though analysis of the genetic control of the response to combined stresses is rare, we have more information about physiological responses for combinations within one or a few genotypes. For example, plant protection chemical mixtures show interaction effects (Dashevskaya *et al.* 2013), as do biotic/abiotic combinations (Prasch and Sonnewald 2014; Kissoudis *et al.* 2014; Suzuki *et al.* 2014). A common theme across all the types of combinations examined is that the effect of the stress interaction is not easily derived from single-effect responses. It is also clear that application of a single, often severe stress treatment is not predictive of response in lower stress levels (Mittler 2006; Tardieu 2012; Zandalinas *et al.* 2017).

Work on mixtures of toxins illustrates the classes of phenotypes one might see in response to combined stresses. Organismal responses to mixtures of drugs and chemical toxins are grouped into modes of action such as concentration addition/independent action, synergy/antagonism, dose-level, and dose-ratio (Jonker *et al.* 2005). The shape of the responses to the mixtures defines these different modes of interaction. In favorable cases, mechanistic inferences can be drawn from an analysis of the phenotypic responses to increasing levels of abiotic stress. Sigmoidally shaped functions indicate a buffered signaling system, while peaks suggest the system includes a negative feedback that damps the response at higher levels. These inferences can then be incorporated into more complex mathematical and network models (Lucas *et al.* 2011; Keurentjes *et al.* 2013). (Reymond *et al.* 2003). As in the univariable, single stress dose-response case, the more complex interactions of mixtures with the organism require more parameters to fit the observed response to a model. Model complexity penalties are thus needed for testing such fits. Response surfaces generated from more than one input variable are most similar to multi-input analog circuits, and analog systems are increasingly prominent modeling approaches in the natural sciences (Sarpeshkar 2014).

Increased growth and yield in maize under drought and low nitrogen are genetically correlated; selection for one stress results in enhanced performance in the other stress environment (Weber *et al.* 2012). However, the correlation can vary by trait and by the specifics of the stress level and genotype used (Bennett *et al.* 1989; Sadras and Richards 2014). Typically, only a few levels of nitrogen and drought are applied; and factorial analyses are used instead of surface-fitting approaches that could compare equivalent stress intensities. For example, the half maximal effective concentration, EC50, is often used as a comparison point in toxicity studies (Piegorsch and Bailer 2005), but has not been widely used for comparisons of abiotic stress tolerances in plants (Claeys *et al.* 2014). Moreover, the EC50 compares values of a single scalar, rather than the *n*-dimensional surfaces that more accurately characterize the response. To better understand the mechanisms of field-relevant stress responses, the interaction between limited nitrogen and limited water in maize should be examined over a large range of levels of the stresses and in multiple genotypes.

The maize inbred lines B73 and Mo17 have different responses to drought and nitrogen. B73 exhibits top-fire and Mo17 barrenness under drought (A. Hallauer, personal communication), and their response to nitrogen differs by *≈* ≈ 25% (Balko and Russell 1980). These two inbreds are known to combine well as a hybrid (Hallauer *et al.* 2010), with the hybrid having very good performance under drought stress (A. Hallauer, personal communication). We infer that there are interactions between alleles of sets of genes in these parents that confer increased stress tolerance in the hybrid. If this were the case, a population of offspring from these parents would generate a range of allele combinations from those sets of genes and would exhibit different responses to varying stresses. Recombinant inbred populations are especially useful for detecting important alleles that segregate from the parent inbreds. Intermating of early generations of offspring to produce recombinant inbred lines (RILs) increases detection power (Lee *et al.* 2002). For labor-intensive phenotype scoring and scoring of the same genotype in many environments, intermated RIL populations optimize power to detect loci, though they do not provide single-gene resolution.

To date there is little information on the genetic architecture of differences in response surfaces. Identifying models that describe phenotypic responses and the alleles that control model parameters is helpful in optimizing crop improvement strategies. In crops this has been carried out for single stresses such as drought, but there is little information on abiotic stress mixtures. As drought and limited nitrogen are the two most critical stresses for maize production, and we already know that these two stresses interact, combining them in a response surface design is a logical next step. We infer from stress combination experiments and toxicological data that sets of specific alleles should be able to control sensitivity of growth to combinations of stress signals — not just the perception of such signals — and thus shift the stress-tolerance system in maize among more-responsive and less-responsive states. We detect such genetically controlled differences in response state by mapping the maize alleles that control this shift in the response surfaces to increasing abiotic stress across a range of stress combinations. The experimentally observed response surfaces delimit parts of a larger phenotypic space. We model points in this space corresponding to the experimentally observed phenotypes with a nonlinear producing function that combines two functions that depend on both stresses.

## Materials and Methods

### Seed Stocks

The *Zea mays* intermated recombinant line population (IBM94) derived from inbreds B73 and Mo17 (Lee *et al.* 2002) was provided by the Maize Co-op (**http://maizecoop.cropsci.uiuc.edu**). Seed stocks were increased using standard nursery conditions at the North Carolina Central Agricultural Station, Clayton, NC. Seed lots were genotyped using eight simple sequence repeat markers; lines Mo066 and Mo062 failed genotyping quality control and were thus removed from the data analysis. The B73 parent inbred was used for random checks across factor levels within the experiment.

### Experimental Design and Plant Growth

A face-centered cubic experimental design (Pukelsheim 2006) with five levels of drought and five levels of nitrogen was used to examine dose response surfaces for mixtures of the two stresses. The statistical program JMP v.6’s (SAS, Inc., Cary, NC, USA) experimental design module was used to compare design matrices and to generate the face-centered cubic sample points (see Supplemental Figure 7). This experimental design has more biological replicates in the center portions of the response surface to enable better fit of nonlinear functions. The experiment was conducted in the Cape Fear Community College horticulture greenhouse (GPS coordinates Lat: N 34*^◦^* 19′ 24″ (34.324*^◦^*) Lon: W 77*^°^* 52*^0^* 45*^00^* (-77.879*^°^*), weather station KNCCASTL2) from May–July 2011. Greenhouse maximum temperature was set to 38*^°^*C.

Slow-release fertilizer was custom-mixed by Coor Farm Supply, Smithfield, NC, with clay pellets containing standard trace minerals, 15% potassium, 15% phosphate, and nitrogen levels of 0, 5, 7.5, 12.5, and 15%. For each fertilizer treatment level, 6.36 kg of fertilizer pellets were mixed with a 0.08 m^3^ bag of MetroMix360 potting mix (SunGro, Vancouver, BC, CA). Deep plant pots (MT38, 0.9 l, Stuewe and Sons, Tangent, OR, USA) were filled with soil-fertilizer mix. Random soil-filled pots were weighed, with an average weight of 350 g per pot. Seeds were planted 1 cm below the soil surface. A random number was generated for each plant pot within each water level using SAS v9.2 (SAS Inc. Cary, NC, USA). The plant pots were sorted by random number within each water level, so that neighboring plants were of randomly chosen genotypes and nitrogen levels. Water evaporation in plant pots containing B73 checks in the greenhouse was examined May 20–24; the average difference in weight between fully wet and dry pots over 24 hour was 170 g. Drought was applied to experimental groups using this average, with drought levels of 8, 20, 50, 80, and 92% of full weight (13 ml water, 34 ml water, 85 ml water, 136 ml water, and 156 ml water applied *per* day *per* pot). Selective watering in different amounts was applied from 20–30 days after planting, beginning when the check plants were at the four-leaf growth stage. Soil water potential was measured with a conductivity meter (EC-5, Decagon Devices, Pullman, WA, USA); the selective watering was stopped when the B73 check 8%-weight plant pots had an average water potential of 2 percent. All pots were watered fully for five days after drought treatment.

### Trait Data Collection

Each plant pot was photographed against a 1 cm grid background 14 days after planting, before selective watering. Plants were re-photographed using the same setup and focal length 35 days after planting, after recovery from selective watering.

Plant photographs were measured using ImageJ (Schneider *et al.* 2012), with the internal centimeter ruler in each image used to calibrate the pixel lengths for each measurement session. Each person analyzing the images practiced on a calibration image set until his or her accuracy was greater than 95%. All plant images are available upon request. In our cubic-centered face experimental design, the maximum sample size was either *n* = 4 or *n* = 8. The complete trait data file is included as Supplemental Data Files 2 and 3.

### Parental Inbred Analysis

The initial plant height was thus subtracted from the final height to generate *z*, the growth variable difference in height. Measured initial, final and difference in plant heights for each parental inbred for each stress treatment were fit with a full factorial model (see Equation 1) using JMPv11 (SAS Inc., Cary, NC).

### Mixture Surface Parameter Fits

For the check B73 inbred with adequate data points, the height difference data were analyzed by the procedure of Jonker *et al.* (2005) using Excel Mixtox analysis tools provided by C. Svendsen (**http://www.ceh.ac.uk/products/stats/mixture-toxicity-analysistools.html**). The Mo17 data had too few points to generate a fit. The first step in the Mixtox analysis procedure was to fit a single dose response relationship to the height difference data using the log-logistic two-dimensional surface^1^ as a dose response model to determine the separate, single parameter effects of water and nitrogen. To fit the surface to the water level, the data were filtered to only include points with the lowest level of nitrogen that had a variation in the water level — in this case, the nitrogen level was held constant at 2.5% in order to analyze the single parameter effect of water. Similarly, the surface was fitted for single-parameter nitrogen by holding the water level constant at 20%. These fitted surfaces are the empirical analogues of the partial derivatives of the response surface with respect to water or nitrogen, constrained to lie on the planes where nitrogen = 2.5% and water = 20%, respectively. The fits of the log-logistic surfaces were further refined using the Solver add-in in Excel to minimize the sum of squares (SS) between the actual data points and the predicted model values. The second step in the analysis was to fit the mixtox mixture dose-ratio and dose-level reference models and the deviation models to the data. In order to optimize use of the solver, which was set up use one measurement rather than replicates and was not optimized for a cubic centered face design, an average was taken for each of the treatment combinations in the larger data set.

### QTL Analysis of Quadratic Fit

As seeds germinate at different times, all the recombinant inbred analyses were conducted on trait measurements adjusted for initial plant size. To determine which markers are responsible for creating significantly different dose response surfaces, we fitted the response data for each marker to a smoothed surface, then compared these to the surface fitted to all the markers from all lines using an F-test (see Supplemental Materials 6). Since our experimental design was optimized to detect interactions among markers and stresses, we fit the data to a quadratic function; and we focused on smooth surfaces to incorporate all the information across levels efficiently. The smoothed surface used to fit the data is Equation 1:

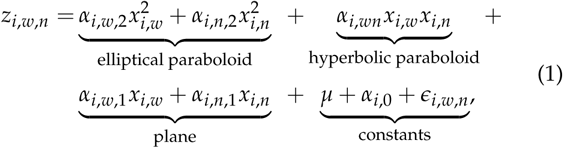

where for each inbred line *i* treated with *x_i,w_* amount of water and *x_i,n_* amount of nitrogen, *z_i,w,n_* is the observed phenotype of that line for that combination of water and nitrogen; *μ* is the mean; *α_i_* is the random effect of the line; and *ɛ_i,w,n_* is the residual error in the fit. The equation is the sum of four component surfaces, marked underneath it.

First, an all-inclusive model was created that included all the markers from all the lines (Motulsky and Christopoulos 2004). Then for each marker, data from all the lines and all combinations of water and nitrogen were divided into two groups according to whether the genotype of the marker was B73 or Mo17. The SAS procedure PROC MIXED was used to model each surface. Due to non-random relatedness between recombinant inbred lines, we incorporated kinship matrix information into the analysis, as recommended by Malosetti *et al.* (2011); kinship matrices were calculated using the SPAGEDI method (Hardy and Vekemans 2002) within the TASSEL v3 program (Bradbury *et al.* 2007).

The sums of squares for the individual marker models were compared to the sums of squares of the all-inclusive model *via* an F-statistic. The resulting raw *P*-values from the analysis were adjusted using the approach described by Makumburage and Stapleton (2011), by grouping correlated adjacent marker *P* values with the Simes function in SAS (PSMOOTH). SAS data steps were used to scan the Simes-adjusted *P* values for groups of significant markers adjacent along the chromosome. A false discovery rate of 0.05 and a Sidak adjustment of 0.05 were each separately used to adjust for multiple testing. Marker data for each mapping line, SAS code for surface fits, *P*-value adjustment, and raw and adjusted *P*-values are provided in the Supplemental Data and Methods files (4, 5, and 1).

### Response Surfaces of the Markers

The fitted response surfaces were plotted in three-dimensional Cartesian space using R, recentering the intervals for water and nitrogen (plotting details and code are provided in Supplemental Methods File 11). For each marker, two response surfaces were plotted, one for the B73 and the other for the Mo17 alleles in the QTL region.

### *Annotation* of QTL *Loci*

QTeller (**http://www.qteller.com**) was used to assemble a list of maize genes in the three chromosomal regions containing QTL that changed the difference in height *z*, and the gene IDs were placed into AGRIGO (Du *et al.* 2010) for annotation analysis. All GO annotations within each QTL were used to create scaled semantic-similarity plots through the Revigo interface (Supek *et al.* 2011).

### Modeling the B73 Surface with a Producing Function

The experimental response surfaces are individual points in a higher-dimensional phenotypic space. This space represents all possible combined stress phenotypes, including ones not yet observed. Thus, exploring that space can identify novel, feasible targets for crop improvement and provide mechanistic insight into the stress response. While Equation 1 suffices for the statistical fit of the data, it is less well suited to exploring the variety of stress responses. First, it offers little insight into possible mechanisms of the responses. Structurally, the equation includes both the elliptical and hyperbolic paraboloids, two “faces”, or complex conjugates, of a larger function that subsumes both, with little to suggest why one paraboloid might be favored over the other in a particular phenotype. Second, of its eight parameters, much of the numerical effect in the fits is carried by the constants that shift the surfaces along the *z* axis (see Table 1). These shifts are extraneous to the surfaces, and beg the question of what drives their values. Under these circumstances, it would be too easy to simulate phenotypes that had little intrinsic relationship to plant’s physiology. We therefore sought a simpler expression that could reproduce the shape of the response surfaces to a reasonable approximation.

The single global maximum of the B73 surface immediately suggested that the simplest function modeling the network responsible for the phenotype would involve a function that generates a peak. The simplest way to produce Mo17’s trough would be to “flip” that peak with a mathematical operation that seems plausible. The sharpness of the B73 peak immediately suggested an elliptical paraboloid. A top-down (orthogonal) projection of such a paraboloid’s peak into the water-nitrogen plane gives an ellipse. For the response surfaces, the ratio of the major and minor axes of the ellipses reflect the relative weighting of water and nitrogen input variables for that phenotype. An elliptical paraboloid is the simplest function that generates a single, easily flipped peak. A two-dimensional Gaussian function is structurally more elaborate and flipping requires the reciprocal of the Gaussian. Periodic functions, such as transcendental or Bessel functions, would have forced us to assume either that their other peaks lay outside the evaluation interval or that the functions were severely damped.

However, two asymmetries in the response surface indicate the producing function is not just an elliptical paraboloid or a similar peak-producing function. First, the maximum is not centered at (0, 0, *z*), but is shifted in the water-nitrogen plane. Second, the surface is “tilted” in the space so that the surfaces formed by the intersection of the surface with the planes bounding the evaluation intervals (*−x_w_z*, *x_w_z*, *−x_n_z*, and *x_n_z*) are not identical to each other. While the peak can be shifted by adding a constant along *x_w_*, *x_n_*, or both, tilting requires the addition of another function. We experimented with many possibilities for this second function and operators to combine the first and second functions, but found the simplest was to add a plane (see Equation 3). The function was computed over the recentered water and nitrogen intervals [*−*42, 42] and [*−*7.5, 7.5], respectively, forming a 169 *×* 31 matrix *z* of 5,239 cells whose values specified the shape and position of the phenotypic surface. We tested candidate combinations of the two functions by deleting equation terms (which can be interpreted as network nodes) and by varying the values of the parameters to confirm our hypothesis. Deletion analysis was also recently recommended as good practice for systems modeling (Babtie *et al.* 2014). Code to generate the surfaces of candidate functions was implemented in R (Supplementary File 12). Surfaces were plotted using the R package rgl (Adler *et al.* 2017–present) in a standard orientation (Supplementary File 16).

### Generating Parameter Values for the Producing Function using Linear Models

To ask how well the producing function (Equation 3) accounts for the smoothed experimental surfaces, we fitted parameter values for using two different linear solvers^2^. We set the value of *c* to *{−*1, 1*}* for the domed and trough-shaped experimental surfaces, respectively, and estimated the values of (*a*, *b*, *d*, *e*) for

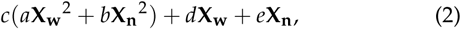

where *X_{n,w}_* are the vectors of *x_w_* and *x_n_* at sets of mesh points, respectively.

The mesh points for these fits were defined by the intersections of surface axes and contours. For the domed surfaces, a central axis was defined as the apparent major axis of the distorted ellipsoid. For Mo17, the central axis was defined as midway between the asymptotes of the trough. For both types of surfaces, rays extending from the peaks at fixed angles relative to the central axes were defined, and the intersections of those rays with a set of contours at fixed *z* values defined the mesh points. These sets of mesh points were then passed to the linear solvers. We used the linear solvers `Solve` and `lsei` in the R package `limSolve` (Soetaert *et al.* 2009; Van den Meersche *et al.* 2009). Both gave identical parameter values for these mesh points. We report those obtained by lsei, since that algorithm also computes the square root of the least square error fit. We repeated these computations using the relative mesh points described in the next section, but the fits were considerably worse except for Mo17, as judged by the cumulated error measures. R code for these computations is in Supplementary File 15.

### Testing the Fit of the Producing Function to the Experimental Surfaces

We next tested whether Equation 3 could reproduce the smoothed, experimentally determined surfaces using the parameters obtained by the linear fits. The most distinctive and robust features of these surfaces are their shape and approximate position in space, rather than their precise location. Because varying model parameters can change both shape and the position of the surfaces in three dimensions, we are interested in shapes defined relative to the peak of each surface, rather than with respect to absolute coordinate values. The mesh points defined above do not capture this shape information well. They are very sensitive to rotation of the simulated surface relative to the experimental, displacement of the surface along the *z* axis, and dilation and contraction of the surfaces’ shapes.

Instead, we defined a set of mesh points that better capture the intrinsic shapes of the surfaces. The peak of each surface lies near a corner of the water-nitrogen (*x_w_*, *x_n_*) plane, so we defined a crow’s foot mesh that radiates away from the peak and towards the two opposite edges of the plane in two steps. First, we defined a set of rays radiating from the peak at fixed slopes relative to the origin, which is in the center of the *x_w_*, *x_n_* plane. plane. Moving the peak relative to the *x_w_*, *x_n_* plane changes the region of the surface spanned, the length of the ray segments, and the absolute position of the mesh points projected onto the evaluation plane. Domed surfaces (those other than Mo17) had rays at slopes (0, 0.05, 0.099, 0.175, 0.32, 0.75) relative to the origin. For Mo17, the slopes were (0, *−*0.05, *−*0.099, *−*0.175, *−*0.32, *−*0.75). Second, we placed 10 mesh points along each ray at the same relative distance from the peak, so 0.1 of the length of each ray segment lying between the peak and the edges of the evaluation *x_w_*, *x_n_*

To compare the smoothed experimental surfaces to those generated from the fitted parameter values, we computed three parameters for each pair of corresponding mesh points:

- *ρ*, the signed Euclidean distances between the surfaces;
- *θ*, the rotation between the experimental and simulated surfaces, projected into the *x_w_*, *x_n_* plane; and
- *δz_r_*, the discrete difference of the relative changes in *z* along the rays.

*ρ* estimates the displacement of the simulated surface in *x_w_*, *x_n_*, *z* space relative to the experimental, due to either or both components of the model. *θ* accounts for different amounts of rotation over the surfaces, due to tilting of the planar component of the producing function. *δz_r_* captures differences in the “bending” of the surfaces, due to either or both components of the model. All three parameters allow for intersecting surfaces. We computed these parameters using all possible pairs of experimental and simulated surfaces. Simulation of Mo17 using the fitted parameters placed the maximum in the corner of the plane opposite that of the experimental surface. Similarly, simulation of QTL3-Mo17 using the fitted parameters placed the peak at the lowest edge of the evaluation plane. These errors in peak placement made computation of *ρ*, *θ*, and *δz_r_* moot. *R* code to generate mesh points, compute comparison parameters, and plot planar projections and heatmaps is in Supplementary File 14. We used the R package superheat for the heatmaps and the image function of package graphics for the planar projections (Barter 2017–present; R Development Core Team *et al.* 2017–present). Both plots used the viridis package for color maps that are less problematic for those with color blindness (Garnier *et al.* 2017–present).

### Data and Code Availability

All supplemental files (input data, SAS analysis code, outputs, supplemental methods, supplemental results, and outputs are available from FigShare at **https://figshare.com/s/3ef69b44d24d0953d625**. Code for modelling, fits, and simulation is on GitHub at **https://github.com/tonikazic/univariate_dose_response.git** in a public repository.

**Figure 1.**
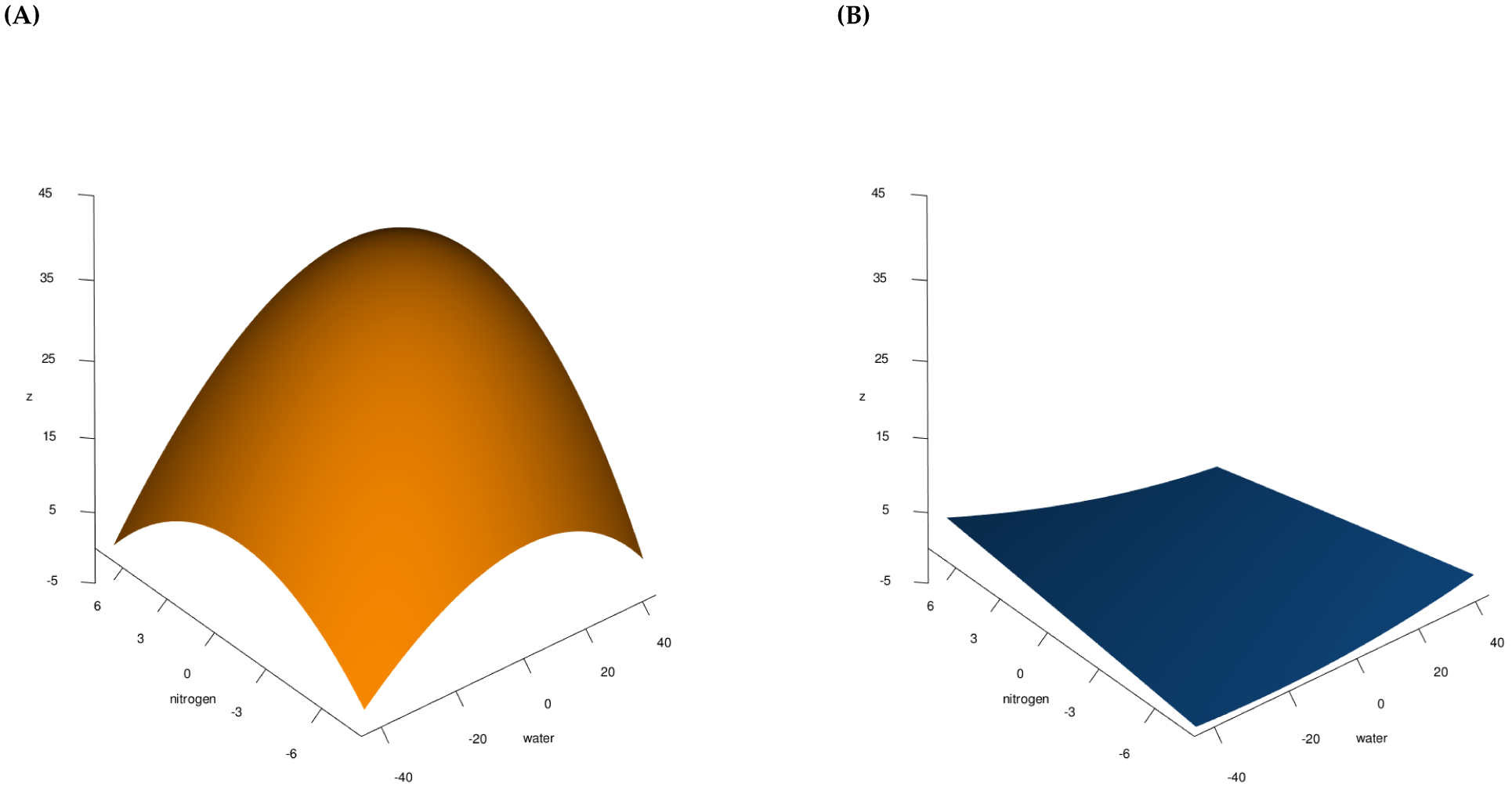
*Parental Inbred Response Surfaces to Combined Drought and Nitrogen Deprivation.* A quadratic surface was fit to the measured trait of differences in plant height (*z*) and is shown for each parental inbred on the same scales. (A) The B73 inbred response surface; (B) the Mo17 response surface.

## Results

### Effect of Stress Combinations on Parental Inbreds

The parental inbreds exhibited different responses to combinations of nitrogen and water deprivation, with B73 showing more variation in its response surface than Mo17 (Figure 1). The plots are rotated to show the most severe stresses in the front center corner, placing the normal full-water and full-nitrogen combination in the back. The intersections of the surface with the height-nitrogen and height-water planes define the phenotypic response at constant amounts of water and nitrogen, respectively, corresponding to the partial derivatives *δz*/*δx_n_* and *δz*/*δx_w_*. B73 exhibited a domed surface, with the peak at moderate amounts of water and nitrogen (Figure 1A). The domed response surface for B73 is convex upward in the sense that it opens downward towards negative values of *z*; and has a relatively high, and highly curved, peak (large *z*_max_ and small discrete curvature, K, that lies in the region of relatively high nitrogen and water (Sullivan 2006). While B73 declined under the most severe conditions (front center corner), it showed modest growth under both moderate drought and very low nitrogen (*e. g.*, the maximum of *δz*/*δx_w_* at the surface’s right edge) and high drought and middle nitrogen (*e. g.*, the maximum of *δz*/*δx_n_* at the surface’s left edge). The worst condition was minimum water and maximum nitrogen (rear left corner).

In contrast, Mo17 had little growth change under any stress (Figure 1B). Its surface is a very shallow trough, or is concave upward; *|z*_min_*|* is small; 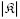 is larger; and the peak lies on a corner. For Mo17, reducing nitrogen affected growth slightly more severely than reducing water (compare the slopes along the left front and right front faces of Figure 1B). We investigated whether this small difference in Mo17’s change of heights could be due to its decreased overall growth. We compared the initial plant heights of B73 and Mo17 and found no significant differ ences between them (*P* = 0.18). When we compared the inbred plant heights after deprivation, we found that the B73 plants had more growth and greater differences between treatments than Mo17 (comparing the factors inbred, nitrogen level, water level and inbred by nitrogen level, at *P <* 0.05) (the numerous sample sizes and confidence intervals are reported in Supplemental File 9). But when we scaled the differences in height to adjust for the smaller Mo17 plants at the beginning of the experiment, plant growth during the experiment was more pronounced for Mo17 in mid-level nitrogen and low water-level treatment combinations, while B73 growth was typically greater when more water was available. This indicates that the Mo17 inbred line is less sensitive to drought provided at least some nitrogen was present. Comparisons at very low nitrogen levels did not exhibit any trend toward differences between parental inbreds (Supplemental Figure 8). For both B73 and Mo17, the slopes of the four lines intersecting each pair of the surfaces’ corners are different, indicating the plants’ responses vary with extremal stress combinations.

Mixture toxicity models with two shape parameters, a and b for water and nitrogen, were used to analyze the shape differences in the parent B73 inbred. (These two parameters are *not* the same as the *a* and *b* of Equation 3, and we have typeset them in a different font than is normally used in mixture toxicity papers to emphasize this distinction.) B73 had the best fit to a dose-ratio surface. a = 0.01 and b = 14.81, indicating that the antagonistic effect of combined stresses is caused mainly by nitrogen deprivation. This is consistent with the curvature of the partial derivatives at the edges of the B73 response surface in Figure 1A: *δz*/*δx_w_* is more sharply curved and has a higher local maximum than *δz*/*δx_n_*.

**Figure 2.**
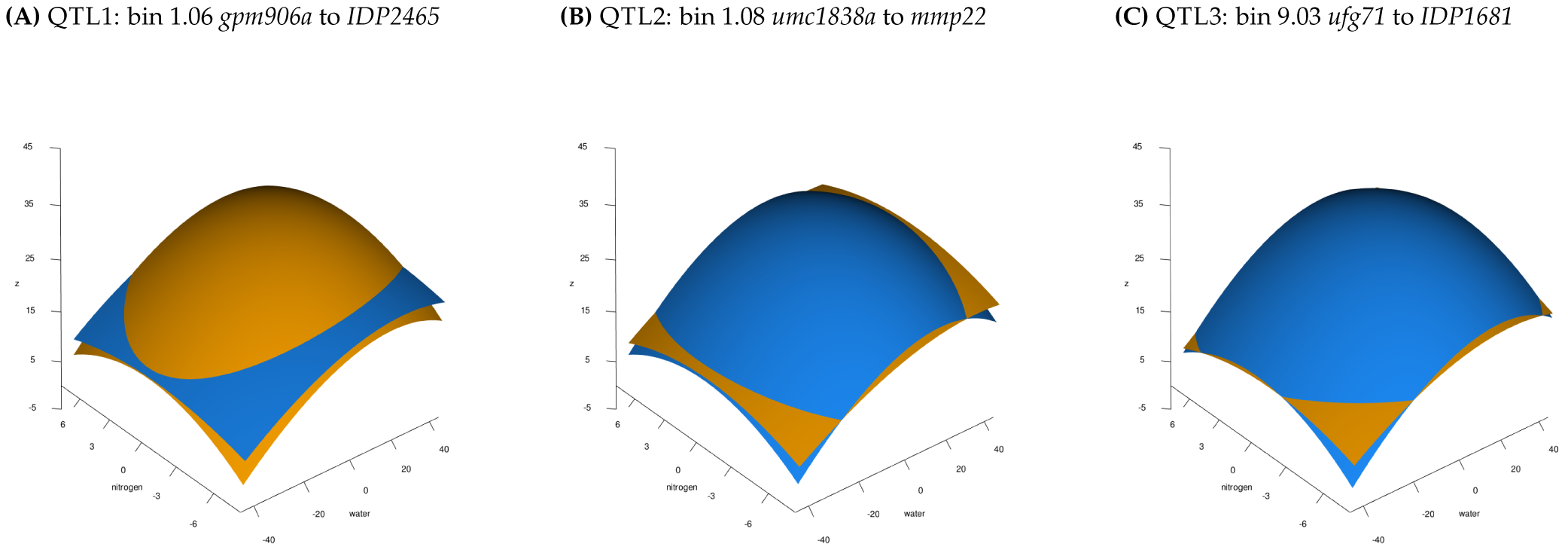
*Loci with Significant Effects on the Phenotypic Response Surface.* Three QTL with a Sidak-adjusted significant interaction surface for differences in height were identified. At each QTL, the trait data were fit with a quadratic response surface separately for each allele. The B73 allele is shown in orange and the Mo17 allele in blue. Each panel shows the QTL’s location (chromosome bin and marker range indicated above) and surface fits for the allele differences at the locus. (A) Surfaces for QTL1, *gpm906a – IDP2465*; (B) surfaces for QTL2, *umc1838a – mmp22*; and (C) surfaces for QTL3, *ufg71 – IDP1681*.

### QTL *that Change Phenotypic Response Surfaces*

Chromosomal loci with significant interaction *P* values for the response surface were fit to nitrogen deprivation and drought, as indicated by intersecting surfaces in the illustrative plots in Figure 2. We found three highly significant QTL that changed the response surface fitted from the data on all lines and an additional 50 QTL that have a false discovery rate-adjusted *P-*values less than 0.05 (Supplemental Results File 9). The three QTL with *P* values below the experiment-wise Sidak-adjusted significance threshold of 0.05 are shown in Figure 2. Table 1 shows the parameter values obtained by fitting Equation 1 to the experimental data.

For all three QTL, the response surface for the Mo17 allele is upwardly convex, a shape resembling that of the B73 allele and parent and very different from the upwardly concave shape of the Mo17 parental surface. In contrast, the response surface for all three QTL’s B73 allele resembled B73’s. The three QTL’s alleles differ from each other and from B73 in many details, including the magnitudes of *z* over their surfaces, the relative magnitudes of the B73 and Mo17 surfaces, and the value and position of *z*_max_. The Mo17 allele’s surface for QTL1, shown in Figure 2A, is pushed upward along the *z* axis, far above the range of the Mo17 parent’s response in Figure 1B. For extremal combinations of water and nitrogen, changes in the growth of the Mo17 allele exceed those for the B73 allele. QTL2’s and QTL3’s Mo17 surfaces lie mostly above those of their B73 alleles; for these loci, the B73 alleles exhibit better performance under extremal conditions (see Figures 2B and 2C). Thus, the phenotypic response surface can differ in shape and magnitude within the population, and surface shape can be quite different than the parent response surface in offspring carrying some QTL allele combinations. Because of the quadratic equation we fit (Equation 1), and the experimental design, which was optimized to detect nonlinear interactions using this quadratic fit, these QTL will have a crossover interaction between the allele surfaces. We show this as intersections between B73 and Mo17 alleles’ response surfaces in plots of the QTL effect (Figure 2). Like their parents, none of these alleles have surface corners that lie on parallel lines.

The differences in surface shape between allele fits in all three QTL, with flatter surfaces with increased combined stress and peaks in the center of the response surface, indicate that nitrogen and water stress have nonlinear effects on plant growth. When these two stresses increase, the combined effect on changes in plant height is less than expected from the independent action of each stress. The domed surface shapes indicate that the better-performing allele at mid-range combined stress is typically not the allele that provides best performance in extreme conditions. The highest water and nitrogen input conditions, which might naïvely be assumed to support the most growth, exhibit less growth and could be favored by a different allele than the mid-range combinations.

### *Annotations of Gene Function in* QTL *Regions*

Gene annotations under QTL provide a qualitative new data type that can provide additional context to the mapping of chromosomal loci. Annotations such as “response to abiotic stress” in the two QTL on Chromosome 1 (Figure 3A and 3B) are consistent with our identification of these QTL as important for response to drought and nitrogen fertilizer. QTL3 in bin 9.03 does not have unique annotations in stress response (Figure 3C); this may indicate that a novel gene type is responsible for the causal allele difference at this locus. The marker with the smallest *P* value within the QTL1 region was *IDP168*, which tags gene *GRMZM5G828396*. This gene is annotated as a basic Helix-Loop-Helix (BHLH) transcription factor. The marker with the lowest *P* value in the second QTL interval was *umc1446*, which tags gene *GRMZM2G162508*; this gene is annotated as a polyketidesynthase-like protein. The marker with the smallest *P* value in the third QTL in bin 9.03, was *mmp17b*, which is between the genes *GRMZM2G538859* and *GRMZM2G093187*; neither gene model has assigned annotations.

### Modeling the Response Surfaces with a Producing Function

We define a complex phenotype as a function of at least three variables, of which at least one is an input, or independent, variable; and at least one is an observable output, or dependent, variable. Here, the phenotype has two input variables, water and nitrogen; and an output variable, relative change in plant height. We assume that all of the observed responses of the recombinant lines are produced by a single network of the organism. Together, these response surfaces form a space of phenotypes. Different surfaces — different points in the phenotypic space — result from different tunings of the network, rather than from fundamental changes to the network’s topology. The greater the range of observed phenotypes, the more the phenotypic space is sampled and the more constraints an hypothesis must satisfy. We call the mathematical function that reproduces the observed phenotypes a “producing function”. The simplest producing function delimits the most parsimonious form of the network, because the two are equivalent conceptualizations of the same biological reality. Tuning the function’s parameters to reproduce the observed phenotypic responses is the same as tuning the network.

**Table 1.** *Parameter Values for Experimental Surfaces Fit to Equation 1.*

| marker | $\alpha_{i,w,2}$ | $\alpha_{i,n,2}$ | $\alpha_{i,wn}$ | $\alpha_{i,w,1}$ | $\alpha_{i,n,1}$ | $\mu + \alpha_{i,0} + \epsilon_{i,w,n}$ |
| --- | --- | --- | --- | --- | --- | --- |
| B73 | -0.006042 | -0.188712 | 0.025567 | 0.229554 | 0.997363 | 32.034927 |
| Mo17 | 0.000814 | -0.001170 | -0.006244 | -0.009502 | 0.063104 | 0.266425 |
| QTL1-B73 | -0.004447 | -0.110527 | 0.007851 | 0.251079 | 0.542153 | 29.572431 |
| QTL1-Mo17 | -0.003192 | -0.049303 | 0.004224 | 0.215691 | 0.317815 | 26.144178 |
| QTL2-B73 | -0.002367 | -0.068065 | 0.006244 | 0.237807 | 0.418609 | 25.779661 |
| QTL2-Mo17 | -0.005370 | -0.105151 | 0.007068 | 0.239113 | 0.497335 | 30.499056 |
| QTL3-B73 | -0.002917 | -0.063999 | 0.008152 | 0.228592 | 0.411333 | 25.720664 |
| QTL3-Mo17 | -0.005307 | -0.116559 | 0.005924 | 0.249475 | 0.525478 | 31.324343 |

**Figure 3.**
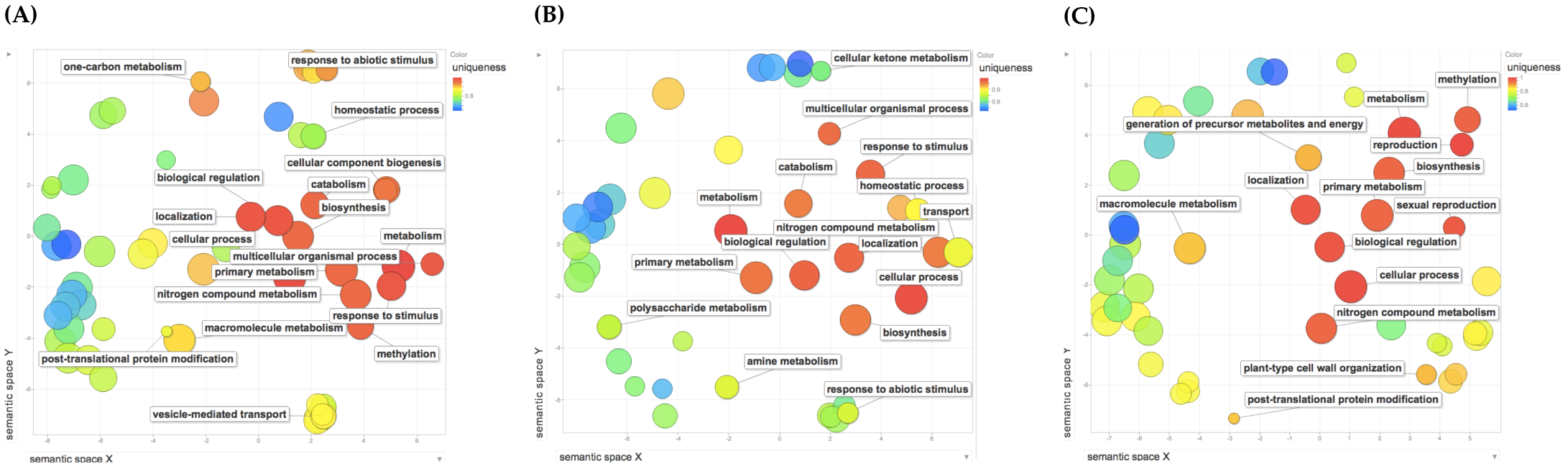
*Gene Ontology Annotations for* QTL. All known genes in each QTL region were scanned for significant annotations. GO process annotations are shown. Annotations were ordered by semantic similarity (Supek *et al.* 2011), with single genes under the QTL having higher uniqueness (more red color). (A) QTL1, bin 1.06, bounded by markers *gpm906a* to *IDP2465*; (B) QTL2, bin 1.08, bounded by *umc1838a* to *mmp22*; and (C) QTL3, bin 9.03, bounded by *ufg71* to *IDP1681*.

So what is the producing function for the network sampled in this experiment? Inspection of the experimental phenotypes of Figures 1 and 2 suggested the simplest producing function is that shown in Equation 3:

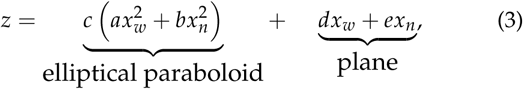

where *z* is the relative difference in height; *x_w_*, water; *x_n_*, nitrogen. *a*, *b*, and *c* are parameters governing the paraboloid’s peak height and orientation (*c*) and its weighting of input water (*a*) and nitrogen (*b*). *d* and *e* tilt the plane along the water and nitrogen axes, respectively. In essence, the model says that the growth response network of maize has two components that read the values of the external water and nitrogen, an elliptic paraboloid and a plane (see Figure 4A).

Both components are essential to reproducing the experimentally observed surfaces, as shown by deleting terms in Equation 3. Omitting the elliptical paraboloid makes it impossible to reproduce any experimental surface, since all have domes or troughs; and omitting the plane makes all of the surfaces symmetric about the major and minor axes of the paraboloid and place the maxima and minima at (0, 0, *z*). The relative weightings of water and nitrogen within each component are different (*a* and *b vs. d* and *e*), and independent of the weightings of the two components. The relative weighting of the paraboloid and planar components is controlled by all five parameters: *c*(*a* + *b*) *> d* + *e* emphasizes the paraboloid nature of the surface. Unlike Equation 1, where the parameters adjust the qualities of the fit of the equation to the observed data, the producing function explicitly treats the parameters as fundamental model components that adjust the organism’s physiological response to input water and nitrogen.

**Figure 4.**
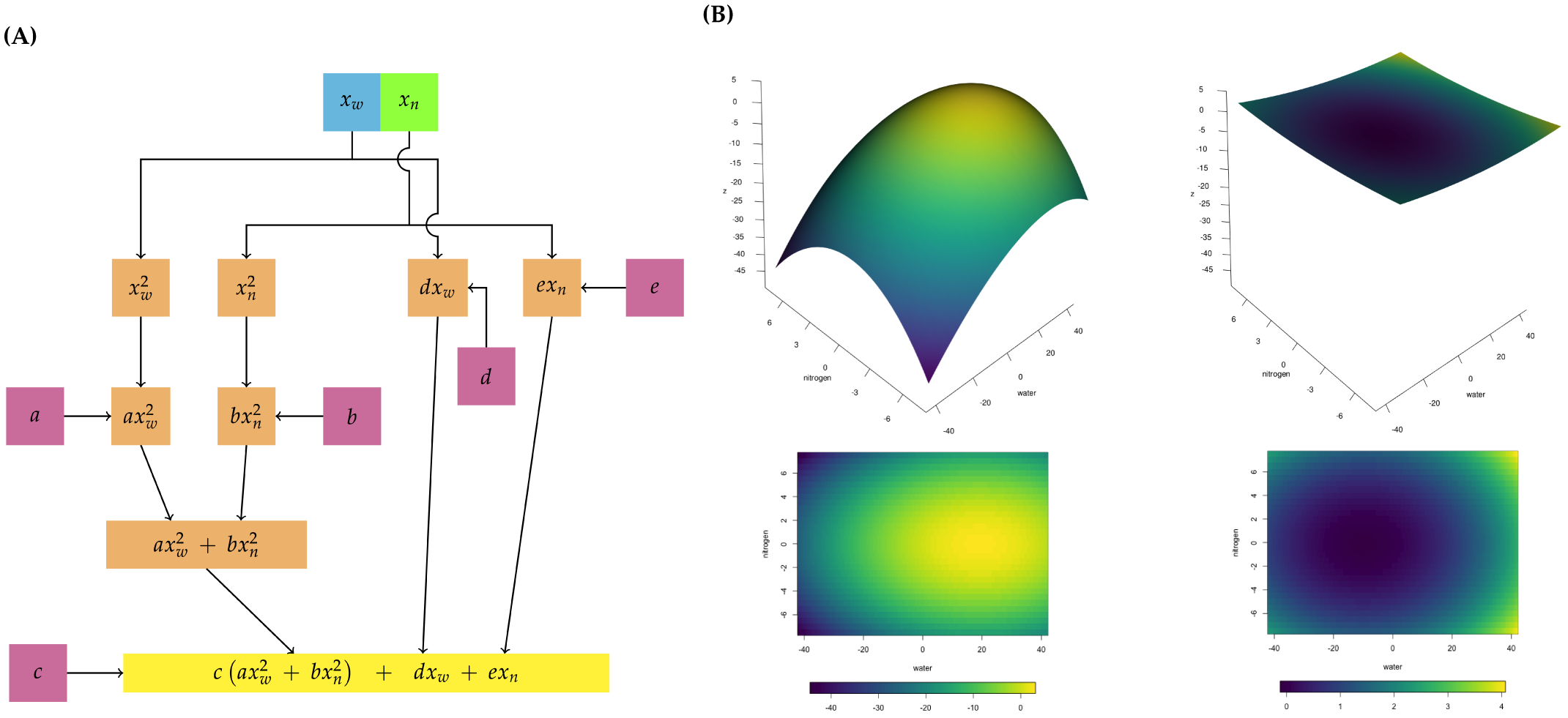
*Modeling of Response Surfaces with the Producing Function (Equation 3).* (A) The producing function drawn as a network. The elliptical paraboloid component is on the left, and the planar component on the right. Each term in the function is a node; the operators are edges. The parameters of the producing function (Equation 3) are in magenta; results of mathematical operations on the input nodes are shown in orange. The final equation is in yellow. (B) Sample simulated response surfaces and their projections into the *x_w_*, *x_n_* plane. (left) B73-like response surface, (*a*, *b*, *c*, *d*, *e*) = (0.260, 0.305, *−*0.175, 2.575, 0.500); (right) Mo17-like response surface, (*a*, *b*, *c*, *d*, *e*) = (0.0330, 0.4250, 0.0030, *−*0.0025, 0.5500). The color scale for each is the same for its surface plot and projection.

The producing function generates the experimentally observed surfaces of the B73 and Mo17 parental inbreds. The simulated surfaces in Figure 4B show good qualitative agreement with the surfaces in Figures 1A and 1B. Setting *c <* 0 makes the surface convex upward, while *c >* 0 produces the concave upward surface of Mo17. *a*, *b*, and *c* together control the depth of the dome or trough. *d* and *e* further adjust the position and magnitude of the surface in all three dimensions, most notably moving the peak about in *x_w_*, *x_n_*, *z*-space. Thus, the phenotypes we observe are products of the entire network.

### The Shapes of the Alleles’ Response Surfaces

How well does Equation 3 reproduce the shapes of the alleles’ response surfaces, the most distinctive and robust feature of the phenotypes? The shapes show the patterns of the alleles’ responses to the stresses, indicate the relative importance of the elliptical paraboloid and the plane, and are less sensitive to the effects of errors due to small sample sizes.

The precision of the reproduction is affected by errors in the data and the nature of the producing function. Optimizing the design of the experiment for peak detection means that the small sample sizes on the edges make it difficult to estimate the surfaces there. As well, the relative “flatness” (small discrete curvature, 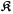) of the peak regions makes the positions of the maxima especially sensitive to difficulties smoothing the experimental data. On the modelling side, the producing function’s planar component cannot reproduce the corners of the surfaces, since these appear to form a nonplanar, twisted quadrilateral (a feature contributed by the hyperbolic interaction term, *α_i,wn_x_i,w_x_i,n_*, of the quadratic Equation 1). Moreover, simulated surfaces that are identical in shape to experimental ones can be rotated in the *x_w_*, *x_n_* plane by minor changes in the values of *a*, *b*, *d*, or *e*, or in the evaluation intervals.

The smoothed experimental surfaces fall into four, *nondisjoint* categories:

**domed** more sharply domed, highest amplitude surfaces with peaks in the high nitrogen, high water region (B73 and QTL1-B73);
**hybrid** higher amplitude domed surfaces with the peaks displaced from the high nitrogen, high water corner towards the center of the water-nitrogen plane (QTL2-Mo17 and QTL3-Mo17);
**shoulder** lower amplitude surfaces, with lower peaks on the high water edge that slope more gradually downward as nitrogen decreases (QTL1-Mo17, QTL2-B73, and QTL3-B73); and
**trough** a very low amplitude trough with a peak at the lowest water and highest nitrogen corner (Mo17).

These categories are illustrated by examining the positions of the peaks in *x_w_*, *x_n_*, *z* (Table 2) and by projecting the surfaces into the water-nitrogen plane (Figure 5). The smoothed experimental surfaces, plotted to show each one’s amplitude, are shown in the first and third columns of Figure 5. We emphasize that the membership of the domed and hybrid categories changes depending on the criteria. If one considers only the position of the peak in the *x_w_*, *x_n_* plane, then B73 and QTL1-Mo17 only would form the domed class, and QTL2-Mo17 and QTL3-Mo17 would form the hybrid class. Weighting the position in *z* more than the peak position classifies B73 as domed and QTL1-B73, QTL2-Mo17, and QTL3-Mo17 as hybrid. Binning the *z*_max_ more coarsely would eliminate the hybrid category all together. Adding other univariate proxy criteria might further change the classification.

**Table 2.** *Coordinates of the Peaks of the Smoothed Experimental Surfaces.*

| proxy | B73 | QTL1-Mo17 | QTL2-Mo17 | QTL3-Mo17 | QTL1-Mo17 | QTL2-B73 | QTL3-B73 | Mo17 |
| --- | --- | --- | --- | --- | --- | --- | --- | --- |
| $x_w$ | 28.5 | 31.5 | 24.0 | 25.0 | 37.0 | 42.0 | 42.0 | -42.0 |
| $x_n$ | 4.5 | 3.5 | 3.0 | 3.0 | 5.0 | 5.0 | 6.0 | 7.5 |
| $z_{\max}$ | 37.6 | 34.5 | 34.2 | 35.2 | 30.9 | 33.3 | 32.4 | 4.5 |

### Comparison of Experimental and Simulated Surfaces

We fitted linear models to a set of points placed on the experimental surfaces that radiated away from the peaks and covered the central, best-determined parts of the surfaces. The points avoided the edges and corners of the surfaces, where the experimental design used the smallest sample sizes. We used models with and without the experimental maxima to see if these affected the fitted values.

Table 3 summarizes the values of *a*, *b*, *d*, and *e* and the cumulative, scalar errors for each smoothed experimental surface obtained by the linear fits using the mesh points. In general, the fitted values distribute the numerical weight more evenly among the four parameters than the fits to the quadratic model of Table 1, even after the lumped linear constant *μ* + *α_i_*_,0_ + *e_i,w,n_* is excluded. The exception is QTL3-Mo17, where nearly all of the numerical weight is concentrated in *e*. Identical parameter values were obtained by both fitting algorithms; the Table shows the lsei results since this method also reports the cumulative scalar errors. The mesh point sets used here included the experimental peak. Omitting these slightly degraded the quality of the fits, as estimated by the cumulative error scalars, for all alleles except Mo17 (Supplemental Figure 10).

We report the square root of the solution norm, which is the minimum of the least squares fit to the equation. The residual norms for all the parameters was 0. We summarize the pattern of parameter signs in the right-hand section of Table 3. The pattern of pluses and minuses groups the surfaces into the two parental types, (domed and trough); flatter, more shouldered surfaces (QTL1-Mo17, QTL2-B73, QTL3-B73); a group of moderately peaked surfaces (QTL1-B73 with QTL2-Mo17) that fall into either the domed or hybrid categories, depending on the weighting of the classification criteria. Finally, QTL3-Mo17 is a hybrid surface based on the peak in *x_w_*, *x_n_*, *z* and the projection into the evaluation plane, but has a distinct sign pattern. Grouping by signs makes it more visually obvious how the surface fits in Figure 5 connect to the parameters in the function, and emphasizes the overlap among the categories.

We used the parameter values from the linear fits shown in Table 3 to generate simulations of the alleles’ response surfaces. Figure 5 shows the smoothed experimental surfaces and their simulations projected into the water-nitrogen plane. These are plotted using the minimum and maximum of each surface to set the scales. The producing function reproduces the major shape features of the experimental surfaces. All four nondisjoint categories of surfaces are generated with the correct membership, and the corresponding differences in their shapes relative to the experimental shapes fall into the same categories. Moreover, the simulations show the expected sensitivities to sample sizes, smoothing problems, lack of twist, and small shifts in fitted parameter values. The fitted values rotate all surfaces counterclockwise and increase their amplitudes. They also displace the peaks of the domed and hybrid surfaces (B73, QTL1-B73, QTL2-Mo17, and QTL3-Mo17) from the high nitrogen, high water corner towards the low-nitrogen edge, passing through the center of the evaluation plane. A similar displacement is seen for the shouldered surfaces, where the shift is toward the high water edge; and Mo17’s surface is shifted to the low nitrogen, high water corner.

The cumulative scalar estimates of error supplied by lsei cannot capture how the quality of the simulations varies over the surfaces. A linear model will necessarily emphasize the highest magnitude values, which in these cases are the peaks, and can exaggerate their impact in the generated surfaces. For each experimental and simulated surface, we computed a set of relative mesh points that better describe the shapes than those used to compute the fitted parameters. These used a set of six rays emanating from the peak at fixed slopes, placing the mesh points at intervals of one tenth the length of the ray segment bounded by the peaks and the water or nitrogen axes. We then compared the shapes for each experimental/simulated pair of surfaces at each corresponding relative mesh point using three parameters: *ρ*, the Euclidean distance; *θ*, the rotation angle between the surfaces when projected into the water-nitrogen plane; and *δz_r_*, the relative discrete difference along each ray, moving away from the peak. For a baseline, we compared each simulated surface (s) to its experimental one (e). The QTL3-Mo17 and Mo17 simulations produced surfaces that were displaced too far in the water-nitrogen plane to be included in the comparison. These values are shown as heatmaps in Figure 6.

In the heatmaps, bluer shades indicate the closest fit for *ρ*; for *θ* and *δz_r_*, the green color indicates the closest fit. The precision of fits was not uniform across any of the surfaces for any parameter, which is visually apparent as blocks of different colors in Figure 6. Different parameters had slightly different blocks, justifying evaluating the fits as three different matrices. Strikingly, the same experimental surface was often fit equally well by multiple simulations, looking over all three parameters. This is consistent with the nondisjoint subspaces we observe in the experimental surfaces, and suggests we did not overfit the parameters using the linear model. Consistent with this, occasionally the self comparisons were not the best fit.

Judged by *ρ*, the surfaces divide into two groups. Experimental surfaces for QTL1-Mo17, QTL2-B73, QTL3-B73, QTL2-Mo17, and QTL3-Mo17 had the most uniform, best fits when compared to their simulations. The first three are shouldered surfaces; the fourth a hybrid surface with the majority sign pattern; and the last is the hybrid surface with the distinctive sign pattern shown in Table 3. In constrast, B73 and QTL1-B73 had the most poorly fitting surfaces when compared to their simulated versions. Both are domed surfaces, with QTL1-B73 also classifiable as a hybrid surface. We could not compare Mo17 to its simulated version, but it fit all the other simulations about equally poorly, as one would expect. The fits varied across the surfaces. In general, fits were better in the central part of the surfaces, which had a higher density of mesh points. For example in *ρ*, there were blocks of better fit in rays r3, r4, and r5 throughout for most comparisons; the fit decayed as we moved away from the peak and towards the edges of the surfaces.

**Figure 5.**
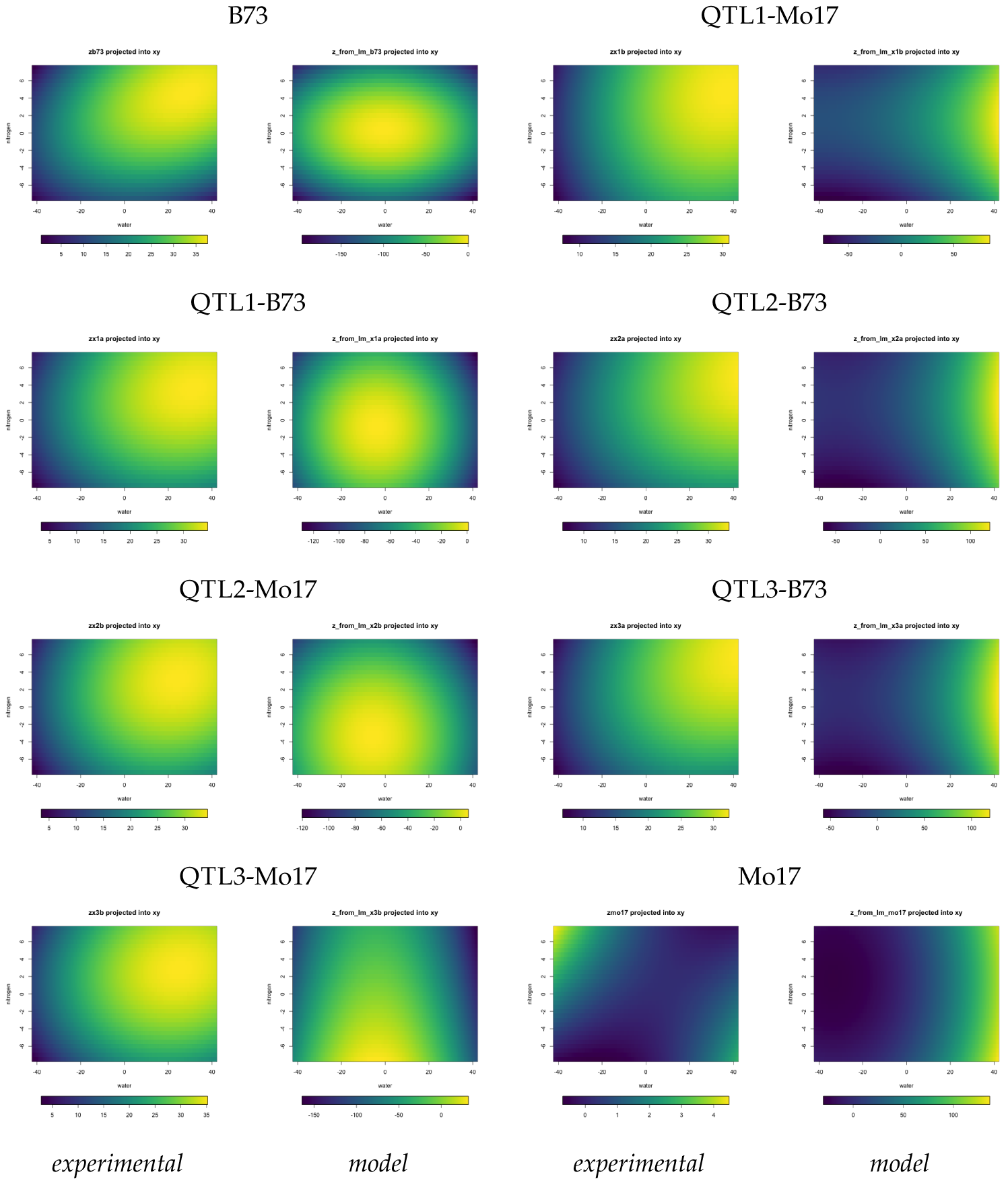
*Experimental and Simulated Surfaces Projected into the Water-Nitrogen Plane.* Each surface’s scale spans its maximum and minimum. The left two columns show the domed and hybrid categories, and the right two columns the shoulder and trough categories. Parameters used in the simulations were obtained from linear fits to the experimental data.

In *θ*, blocks of good quality fit are apparent, and the domed/hybrid, shouldered, and trough sets can be readily identified. As with *ρ*, some surfaces were not always best fit by their simulations; for example eQTL1-B73 to sQTL1-B73 had a poorer fit than eQTL1-B73 to sB73. The rotation of simulated surfaces relative to the experimental surfaces is apparent from the repeating vertical stripes of yellow for ray r1 next to the green stripes for ray r2. The repeating yellow stripes of ray r1 indicate that all the comparisons except those to the experimental Mo17 surface reach their maximum rotation in the upper region of the mesh points. Rotation decays on either side of r1, and is least in the central region of the surfaces. Why? Recall that the rays are of fixed slopes relative to a horizontal ray radiating leftward from the peak. Rotation increases as the difference between the absolute lengths of the rays compared increases: as the peaks from which the rays are drawn shift in the *x_w_*, *x_n_* plane, these ray lengths and the portion of the surfaces they subtend will change correspondingly. Thus, the maximum rotation we observe along ray r1 reflects greater length differences in the r1 rays relative to the other ray pairs. There are two reasons why this could occur. First, *d* and *e* together shift the peaks’ positions and rotate the surfaces, so the rotations would reflect imprecision in the estimates of *d* and *e*. Second, the nondisjointness of the surfaces’ categories may not be intrinsic to the system; perhaps an additional term in the producing function would separate these categories and provide better estimates of all the other parameters, including *d* and *e*, possibly at the cost of more parameter indeterminacy.

**Table 3.** *Parameter Values, Error, and Sign Pattern for Experimental Surfaces Fit to Equation 3.* The experimental mesh points used in the fits included the peak of each surface. The value of *c* was preset to produce the appropriate convexity. The residual norms were all 0.

| *allele* | *a* | *b* | *c* | *d* | *e* | (solutionNorm)<sup>1/2</sup> | sgn(*a*) | sgn(*b*) | sgn(*d*) | sgn(*e*) |
| --- | --- | --- | --- | --- | --- | --- | --- | --- | --- | --- |
| B73 | -0.0463 | -1.8005 | -1 | -0.0765 | 1.3540 | 146.0321 | — | — | — | + |
| QTL1-B73 | -0.0429 | -0.5926 | -1 | -0.3134 | -0.9222 | 127.2874 | — | — | — | — |
| QTL2-Mo17 | -0.0332 | -0.4362 | -1 | -0.3681 | -2.9073 | 134.1696 | — | — | — | — |
| QTL3-Mo17 | -0.0636 | 0.0039 | -1 | -0.5199 | -4.0788 | 112.6761 | — | + | — | — |
| QTL1-Mo17 | 0.0164 | -0.6201 | -1 | 1.2777 | 1.8758 | 89.4613 | + | — | + | + |
| QTL2-B73 | 0.0268 | -0.4119 | -1 | 1.7254 | 1.7119 | 81.4159 | + | — | + | + |
| QTL3-B73 | 0.0278 | -0.4450 | -1 | 1.6600 | 1.0073 | 83.2248 | + | — | + | + |
| Mo17 | 0.0259 | 0.2164 | 1 | 1.7406 | -0.7203 | 34.1571 | + | + | + | — |

**Figure 6.**
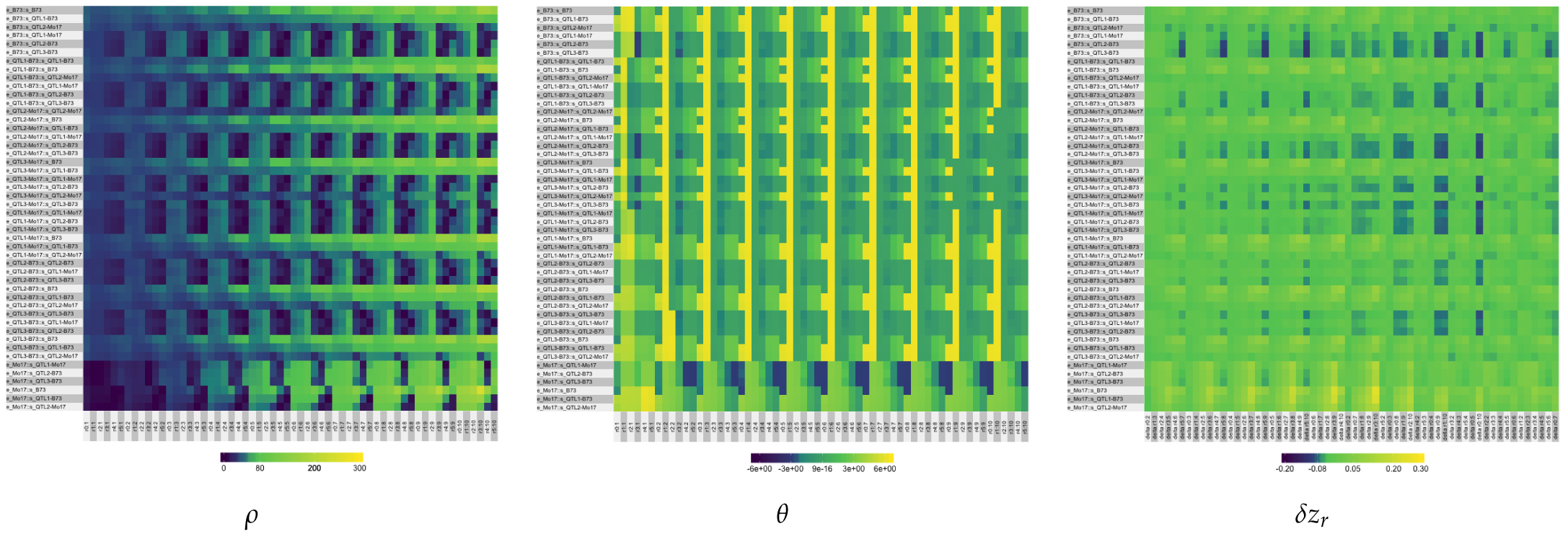
*Heatmaps of ρ*, *θ, and δz_r_*. The pairwise comparison of all 60 relative mesh points for each pair of experimental (e) and simulated surfaces (s) are the rows; the values for the pair of mesh points are the columns. The columns are ordered by concentric rings around the experimental peaks, so that the leftmost six columns are closest to the peak (:1) and the rightmost six furthest away (:10). Within each set of six columns, the rays (r0, r1, …, r5) are arranged in order of increasing slope (non-Mo17) or decreasing slope (Mo17). The scales are heatmap-specific.

The final comparison in Figure 6 is the relative changes in shapes of the surfaces along each ray, subtracting the experimental from the simulated surfaces, *δz_r_*. By this criterion, the shapes are quite similar, as indicated by large areas of green. Ray r0 changes very slowly and uniformly until about mesh point :9 or :10, while ray r5, the most vertical ray, changes fastest and most dramatically. As we descend from the peak, other rays also start to change more rapidly. This is consistent with the changes in *ρ* that we observe: differences in shapes will also appear as differences in Euclidean distances among pairs of points. In this *δz_r_* criterion, the values are negative if the simulation descends more rapidly than the experimental and positive if the surface is shallower. Most changes are towards sharper simulated surfaces (blue), and a few to shallower simulated surfaces (yellow). This is consistent with poorer estimates of the producing function’s parameter values: all four control the sharpness and height of the peak and the shapes of the sides of the surfaces.

## Discussion

The parental inbred patterns of stress response did not predict the responses of the Mo17 QTL alleles. Mo17 is the least responsive of all the lines, and lines with the Mo17 alleles in mixed RIL backgrounds have responses that show much greater, more B73-like amplitudes. The trait response of the Mo17 bin 1.06 QTL allele (QTL1-Mo17) is greater than the corresponding B73 allele. In contrast, the Mo17 alleles for QTLs in bins 1.08 and 9.03 (QTL2-Mo17 and QTL3-Mo17) show more response than their B73 alleles. However in all cases, the response surfaces of the Mo17 alleles are convex upward like the B73 parent. Mo17 appears to contain allele sets that together restrain growth across all environments (Figure 1B). The responses of the B73 alleles are damped and shifted by the addition of Mo17 germplasm in the background. This transgressive segregation pattern is often seen in mapping populations (Rieseberg *et al.* 2003) and indicates that the parental inbreds contained alleles with antagonistic effects on the trait. Transgressive segregation and the changes in response surfaces’ shapes suggest that other modifier genes segregating within the populations are affecting the response to combination stress.

We weighted our statistical model fit for overall polygenic similarity, but specific epistatic interactions using combinatorial methods (Mäki-Tanila and Hill 2014). However, it is likelier that any epistasis one might deduce from a linear model is instead the result of attempting to fit a linear model to a nonlinear response phenotype (Sailer and Harms 2017). A nonlinear function’s matrix of coefficients captures both the interactions among genes and the magnitudes of those interactions. Linear transformations of nonlinear phenotypes partition that matrix into a matrix of interactions — the adjacency matrix of a network — and a matrix of interactions’ magnitudes. The more nonlinear the phenotype, the greater the apparent importance of an epistatic interaction term in the decoupled linear model. Paths through the network select subsets of terms from the full nonlinear model, fragmenting the phenotype into multiple approximations that appear as epistasis. Better nonlinear models would fold these apparent epistases into the inherent nonlinearity of the response, distinguishing true epistatic interactions from nonlinear phenotypes.

Antagonistic, damped responses to combinations of stresses were previously observed in this IBM population in an experiment measuring changes in height under combinations of UV and drought, which identified different QTL than we see here for drought and low nitrogen (Makumburage *et al.* 2013), though different QTL were identified for the UV and drought change in height trait in that work as compared to the QTL we identify in this study for drought and low nitrogen. It is possible that specific stress combinations have different genetic control than the response surfaces that we studied in this work. To test the extent of stress-combination specificity, dose response analyses for more combinations of stresses would be needed. As a first step, before doing many such experiments to attempt to find coincident QTL, we developed a mathematical model for the control of combined stress input and plant growth output.

All three of the QTL we identified have better performance for one allele in the mid-range combined stress, at moderate levels of water and nitrogen. Extreme values are often used in lab-scale studies, though it is difficult to extrapolate from those results to the more common, moderate stress settings (Tardieu 2012). Our QTL suggest that comparison of a control environment with an extreme stress environment could suffer from Type II error, as none of our QTL show a pattern indicative of allelic differences at the most extreme stress levels. This also fits with indirect selection results in hybrids, where optimal conditions show a relatively high level of genetic correlation with stress environment performance (Weber *et al.* 2012). Analysis of selection efficiency on yield test winner plots as compared typical plots, or classification of fields by optimality criteria, may be useful for better prediction of performance in trials. Correlations between optimal and stress environments are typically positive but the extent of the correlation is not always as expected (Weber *et al.* 2012): this could reflect a history of selection for “good-middle” stress performance alleles instead of a selection from a straight linear surface.

More complex surface fits, such as dose-ratio (Jonker *et al.* 2005), provide additional information (and require more data. The B73 MixTox analysis shows that low nitrogen “overshadows” drought; in low nitrogen, having additional water available does not improve growth. The over-riding importance of nitrogen for growth is consistent with agricultural recommendations for modern corn lines (Hallauer *et al.* 2010). Conversely, keeping lower levels of nitrogen fertilization was found to ameliorate the effect of severe drought under certain conditions (Sadras and Richards 2014). Given the transgressive segregation we see in our mapping population, one reason for inconsistency in tests of severe-stress amelioration may be genetic differences in capacity for overshadowing. A comparison of response surfaces in lines selected for stress tolerance compared to unselected lines would be a useful test of the genetic contributions to the interactions between water and nitrogen use.

We observed that the parental B73 and Mo17 show different patterns of combined stress response. These lines were selected to create an excellent single-cross hybrid (Hallauer *et al.* 2010). This suggests that lines with little response or that tolerate extreme stress may combine well with good-middle lines. To test this, relationships between response surface shape to combining ability could be done across a range of good and poor combiner varieties, and across general and specific combiner extremes.

Nonlinear changes in growth in response to combined stresses is a complex phenotype. Just from the surfaces fit to the experimental data, one can rule out an unbranched network sensitive to both water and nitrogen, because the response’s peak does not scale with the sum or product of the inputs. Similarly, one can exclude two completely independent nodes, one for each input. Instead, the two inputs interact so that the response varies as a function of both: the phenotypes are produced by the action of the entire network, rather than just paths through it (Figure 4A). The nonlinear producing function of Equation 3 defines this interaction as the sum of two components, each of which is sensitive to both water and nitrogen. Environmental perturbations, such as varying available water and nitrogen, change the intervals over which the producing function is evaluated by the plant. Genetic perturbations, such as the alleles of the QTL identified in this work, delimit different regions of the space of possible phenotypes. Of course, considering additional phenotypes or phenotypic dimensions might necessitate changing the function.

The producing function is a simpler, more coarsely-grained model that approximates the behavior of the system. It successfully accounts for all the important qualitative features of the experimentally observed phenotypes without assuming large constants of a regression model that here numerically dominate the solution (Table 1). Surfaces generated using parameter values obtained by linear fits approximated the shapes of the experimental surfaces reasonably well, but were shifted downward in three-dimensional space. Slightly tilting the plane component of Equation 3 can strongly shift and rotate the position of the peak in *x_w_*, *x_n_*, *z*. This suggests the model is quite sensitive to variation in *d* and *e* and is consistent with our simulations (data not shown).

Estimating parameters by fitting is more challenging with nonlinear phenomena. Two questions arise: what should be fitted? and how should it be fit? The first question is the fit criterion. We tested a wide variety of objective functions, singly and in combination to produce scalars or vectors, to describe the surfaces through a set of proxies that are easier to compute. None effectively discriminated among somewhat similar surfaces. This is not surprising in retrospect: such a function would behave as a unique, deterministic, multidimensional hash function. This situation is likely to occur when considering complex phenotypes, which are inherently multivariate. Instead, we used three evaluation criteria computed over a set of mesh points spanning portions of the surfaces. Further characterizations of the resulting matrices, instead of visually inspecting the heatmaps of Figure 6, might allow us to condense these descriptions, especially since they exhibit periodic behaviors. Nonetheless, the heatmaps illustrate how much the surfaces can change as a function of the fitted parameter values and suggest a more refined alternative to simple scoring schemes.

The second question is one of fitting method. The most common approach approximates the higher degree polynomial with a linear statistical model, as we have done here. Our results clearly demonstrate how poorly such methods can approximate nonlinear functions. A more sophisticated, but compute-intensive, approach would be to fit a set of planes, approximating the surface as a set of three-dimensional splines. Another approach is to optimize the fit of the model to the data using one of many nonlinear optimization techniques (Bartholomew-Biggs 2008). Nonlinear optimizations require a good initial guess for the parameter values. We found the values generated by the linear fits were not good guesses, evidently placing the initial point outside the feasible region of the optimization. Further experimentation will be needed to improve the fits using this approach.

Finally, one can search systematically for parameter combinations that generate surfaces that match the experimental ones. This is one way of asking how sensitive the phenotypes are to variations in the parameters’ values. For nonlinear polynomial functions, the relationship between sets of parameter values and generated surfaces will not be regular or easily anticipated: stepwise changes in parameter values will produce “clumps” of generated phenotypes. The reasons for this clumpiness are instructive. The producing function maps between the parameter and the phenotypic spaces. This mapping may be bijective (one-to-one and onto): each point in the parameter space identifies a unique response surface — a point in the phenotypic space — and *vice versa*. The most common interpretation of complex phenotypes in QTL experiments is that each is unique and that its components are individually determined. This is consistent with the assumptions that the mapping is bijective; that each component of the phenotype is governed by a single parameter; and that the fundamental properties of the two spaces, such as their classification, point density, and smoothness, are the same.

But our results suggest these assumptions may not always be sound. The phenotypes we observe fall into four overlapping classes: domed surfaces, with the peak somewhat interior in the water-nitrogen plane; hybrid surfaces, with a lower peak shifted further into the water-nitrogen plane; shouldered surfaces, with a still lower peak towards the edge of the plane; and trough surfaces, with the peak at a corner of a very flat trough. The close similarities of the phenotypes within each class, and the overlap between the domed and hybrid classes depending on the classification criterion, suggest that the parameter values governing them can fall into rather broad ranges, a hallmark of sloppiness in model systems that breeders commonly call equifinality and statisticians call “parameter nonidentifiability” (Transtrum *et al.* 2015; Luo *et al.* 2009; Hartung 2014; Hines *et al.* 2014). This interpretation is supported by extensive simulation experiments: so far, we have been unable to identify unique combinations of parameter values that determine each individual phenotype. The parameter ranges for the observed phenotypes are not disjoint, another characteristic of sloppy systems (data not shown). Thus, our data divide both the parameter and the phenotypic spaces into nondisjoint subspaces.

One common suggestion is to map QTL using a producing function’s parameters, rather than using a specific quadratic fit (Reymond *et al.* 2003; Lamsal *et al.* 2017). Here, one asks for alleles that shift the phenotype from one subspace to another, presuming the alleles shift the plant from one parameter subspace to another. The usual objective of QTL experiments is to map a peak that exerts a large change on a single parameter of a phenotype. For example, *c* acts like a repressor in some sense, and so it might seem tempting to fit our trait and genotype data to that parameter in order to map repressor alleles. But the subspaces we have identified are produced by all five parameters, so it is likely that subsets of QTL will influence multiple parameters, perhaps differently; that these QTL subsets will not be mutually disjoint; and the subspaces of the parameters will differ in their stiffness and sloppiness.

We encourage application of our response surface and shape modelling approaches to crop protection mixture or abiotic-biotic mixture experiments to understand if there are a few major-effect or many small-effect genes that control these cases. This would affect the design of multi-environment trials. For modeling and eventual causal understanding we need to know the patterns of phenotypic responses and genetic control: not just specific alleles for specific combinations, but alleles that affect high-level patterns of response, such as the categories of surfaces. For example, single transcription factors are high-level regulatory effectors and have provided effective avenues for crop improvement, and so might be candidates for changing groups of parameter values that then shift higher order patterns. Our work is the first step toward identifying higher order regulatory environmental-response alleles, such as cellular transcription factors and/or physiological regulatory factors such as hormones (Cabello *et al.* 2014). The function we developed for modeling the responses clearly shows the interaction of the two stressors and the range of nonlinear responses of the system. The response surface modelling and shape evaluation methods used here will be useful in detecting such higher order patterns. The overlapping categories formed by these phenotypes may reflect genuine nondisjointness in phenotypic space or missing dimensions that would separate the categories in a higher dimensional space. The methods used here, applied to higher dimensional data, offer one route to distinguish these hypotheses.

## Acknowledgments

We thank Amy Borsay for help with plant handling and data collection in the greenhouse, Maria White for genotyping assistance, Heather Manching and D. Brad Moore for updating our SAS code for grouping and adjusting *P* values, Coor Farm Supply (Smithfield, NC) for donation of custom-mixed fertilizer pellets, and Peter Balint-Kurti, William Beavis, Karen Cone, Nancy Flournoy, Walter Gassmann, Leonard Hearne, Carolyn Lawrence-Dill, Elizabeth Lee, Jonathan Lynch, Mac, Nirav Merchant, Nathan Miller, Martha Narro, Linh Ngô, Avi Vatsa, Vinny, and Ramona Walls for valuable discussions. This project was supported by the National Research Initiative Competitive Grant no. 2009-35100-05028 from the USDA National Institute of Food and Agriculture to AES and by the National Science Foundation’s MCB-1122130 to TK. The funders had no role in study design, data collection and analysis, decision to publish, or preparation of the manuscript.

## Appendix: List of Supplementary Material

- On FigShare at **https://figshare.com/s/3ef69b44d24d0953d625**:

1. Supplemental_Material_metadata_readme.txt
2. Supplemental_Data_File_1a.csv
3. Supplemental_Data_File_1b.csv
4. Supplemental_Data_File_2.csv
5. Supplemental_Data_File_3.csv
6. Supplemental_Methods_File_1.rtf
7. Supplemental_Results_Figure_S1.png
8. Supplemental_Results_Figure_S2.png
9. Supplemental_Results_File_1.csv
10. Supplemental_Results_Figure_S3.pdf
- On GitHub at **https://github.com/tonikazic/univariate-_dose_response.git**, a public repository:
  11. replot MatLab surfaces in R in standard orientation: replot_ann.r
  12. generate surfaces according to Eqn 3: modified_eqn.r
  13. library of proxies and other helper functions: sweep_fcns.r
  14. library of helper functions for analysis: analysis-_fcns.r
  15. linear estimation of parameters: estimate_exptl_parameters.r
  16. standard view for plotting surfaces in 3D: std_view.r

## Footnotes

1 Throughout, we use the more general term “surface” to denote surfaces in any number of dimensions, including the “curves” traced by the successive positions of a single point in an *n*-dimensional space.

2 Throughout the text, we use the term “linear” in its strict mathematical sense when referring to models. The statistical linear models we fitted included quadratic terms (see Equations 1 and 2), in accordance with statisticians’ usage.

